# Evolution of sideways locomotion in crabs

**DOI:** 10.1101/2025.09.20.677458

**Authors:** Junya Taniguchi, Tsubasa Inoue, Kano Kohara, Jung-Fu Huang, Atsushi Hirai, Nobuaki Mizumoto, Fumio Takeshita, Yuuki Kawabata

## Abstract

The evolutionary change in the mode of locomotion is often a major evolutionary event, triggering diversification. Sideways locomotion is a defining feature of true crabs (Brachyura) and may have contributed to their ecological success. Yet, the evolutionary origin of this unique behavior remains unknown. Here we show that the prevalence of sideways locomotion in true crabs reflects a single evolutionary origin from a forward-moving ancestor. Our behavioral analysis of 50 live crab species indicates that crab locomotion can be broadly separated into two predominant modes, sideways and forward locomotion. The phylogenetic comparative analysis revealed a single origin of sideways locomotion, with multiple independent reversions to forward locomotion in ecologically specialized groups. The species richness data show that the lineage in which sideways locomotion originated is far more diverse than its nearest outgroups. These results are consistent with the idea that sideways locomotion acted as a key innovation contributing to the evolutionary diversification of true crabs. Such a rare but innovative behavioral trait provides a framework for understanding how locomotor modes shape evolutionary diversification in animals.

## Introduction

The evolution of novel functional traits can enable organisms to exploit previously inaccessible ecological niches (Stroud & Losos 2016; Miller *et al*. 2023). Such key innovations have shaped biodiversity on Earth by facilitating adaptive radiation within lineages. Because locomotion is a fundamental behavior that is involved in most survival and reproductive processes (Alexander 2003; Domenici 2010), innovations in locomotor mechanisms are often linked to adaptive radiations (Astudillo-Clavijo *et al*. 2015; Higham *et al*. 2015; Burress & Wainwright 2019; Hedrick *et al*. 2020; Feiner *et al*. 2021). For example, the evolution of flight in insects opened aerial niches globally and contributed to their enormous radiation (Grimaldi & Engel 2005), whereas modifications of the locomotor skeleton in *Anolis* lizards facilitated localized adaptive radiations within islands (Feiner *et al*. 2021). Most comparative studies on such innovations in locomotor behavior have taken a functional morphological approach. However, despite the recognized importance of behavioral aspects of key innovations (Miller *et al*. 2023), the actual locomotor behaviors have rarely been compared across species, largely due to the challenges of obtaining large, comparative datasets of animal behaviors.

True crabs (Infraorder Brachyura) are iconic for their sideways locomotion, enabling them to achieve fast bidirectional movements (Vidal-Gadea *et al*. 2008), which may be beneficial for escaping from predators (Wolfe *et al*. 2021). Sideways locomotion is associated with greater joint flexibility in the lateral direction and a thorax elongated along the preferred direction of locomotion (Vidal-Gadea *et al*. 2008). This unique locomotion could be a key innovation in decapod crustaceans, as it changes the behavioral axis through which animals interact with the environment (Miller 1949). Moreover, sideways locomotion could have contributed to the ecological success of true crabs. The number of species of true crabs (∼7,904 species) far exceeds that of their sister group, Anomura (hermit crabs and others; ∼3,437 species), or their closest relatives, Astacidea (clawed lobsters and crayfish; ∼792 species) and Achelata (spiny and slipper lobsters; ∼153 species), according to the World List of Decapoda, DecaNet (DecaNet eds. 2025). True crabs have also successfully colonized diverse habitats globally, including terrestrial, freshwater, and deep-sea environments (Wolfe *et al*. 2024; DecaNet eds. 2025). In addition, the crab-like body plan has evolved repeatedly among decapod crustaceans, a phenomenon known as carcinization (Morrison *et al*. 2002; Tsang *et al*. 2011; Keiler *et al*. 2017; Wolfe *et al*. 2021). Despite this rich diversity and extensive morphological information, however, data on actual locomotor behaviors of crabs are sparse, and no comparative studies based on large datasets have been conducted thus far, making it difficult to evaluate the role of this unique locomotor mode on crab evolution and diversity.

Although most true crabs use sideways locomotion, some groups—including raninids, majids, and mictyrids—move predominantly forward (Sleinis & Silvey 1980; Faulkes 2006; Vidal-Gadea *et al*. 2008). This raises key questions: when did sideways locomotion originate, how many times did it evolve, and how many times did it revert? With a recent comprehensive crab phylogeny based on genomic data (Wolfe *et al*. 2024), here, we conducted behavioral analyses of 50 live crab species. We aimed to (i) pinpoint the origin of the sideways locomotion within Brachyura, (ii) estimate the number of transitions and reversions between sideways and forward locomotion, and (iii) test whether the emergence of sideways locomotion is associated with species diversification. Our results highlight sideways locomotion as a rare but innovative behavioral trait, providing a framework to understand how locomotor modes shape evolutionary diversification in animals.

## Methods

### Experimental procedures

We obtained live crabs from multiple sources, including intertidal and subtidal field collections, public aquaria, and local fish markets. Animals were kept only as long as required to record locomotion and were returned to their habitat or handled according to institutional animal care guidelines. Animal care and experimental procedures were approved by the Animal Care and Use Committee of the Faculty of Fisheries, Nagasaki University (Permit No. NF-0060) in accordance with the Guidelines for Animal Experimentation of the Faculty of Fisheries and the Regulations of the Animal Care and Use Committee of Nagasaki University.

Locomotion was recorded in plastic circular arenas (diameter 80–140 cm) whose medium matched each species’ native environment (dry, seawater, freshwater, or brackish, with or without bare sand) (Fig. 1a). Individuals were acclimated for 5 min in a bucket and then for 1 min inside a transparent cylinder placed at the arena center to minimize startle responses. After removing the cylinder, each trial was filmed for 10 min using a standard video camera (DSC-RX0, Sony Corporation, Tokyo, Japan) at 30 frames s^−1^. The locomotion data were obtained from one representative individual per species due to logistical constraints and the low expected within-species variation. Our preliminary observations on several species for which many individuals were available suggest that locomotor direction is a species-level trait that is typically conserved. Nevertheless, this design does not capture possible ontogenetic, size-dependent, or allometric variation within species. Accordingly, our conclusions are intended to identify broad interspecific patterns of locomotor direction rather than the full range of within-species variation.

**Figure 1.**
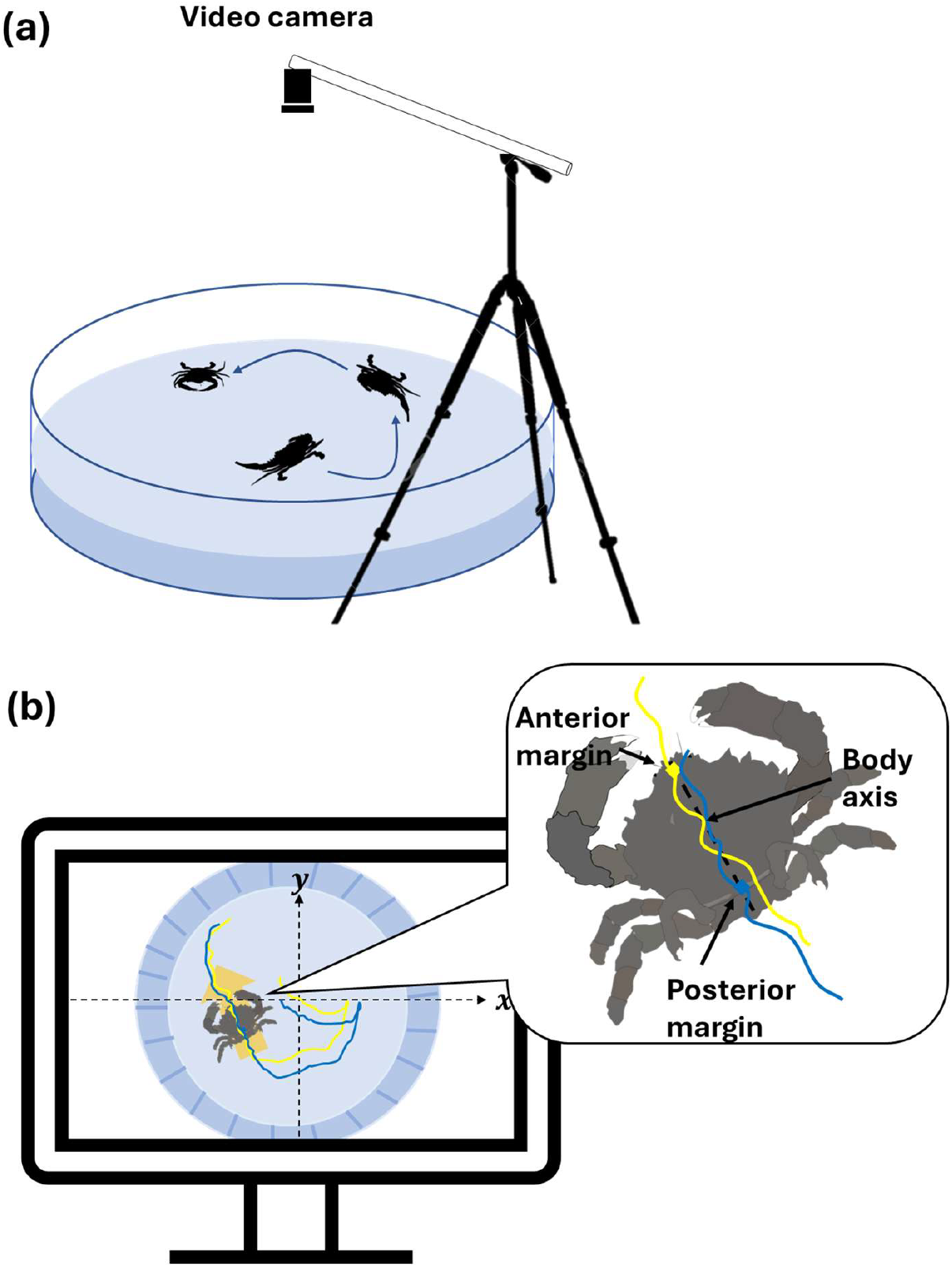
Video acquisition and analysis workflow. (a) Experimental setup used to record each crab’s behavior. (b) Extraction of two-dimensional position coordinates from video frames.

### Video analysis

Obtained videos were converted and downsampled to 5 frames s^−1^ for analysis using XMedia Recode 3.5 (www.xmedia-recode.de). For each frame, two landmarks along the longitudinal body axis (anterior and posterior carapace margins) were digitized using Kinovea 0.8.27 (www.kinovea.org) to estimate the instantaneous body axis and the centroidal position of the animal (Fig. 1b).

To standardize the evaluation of movement directions, we used a reference circle centered on the animal’s starting position (Fig. 2). Each displacement bout was defined as the movement from the starting point to the point where the trajectory of the body center crossed the circle, with the next bout beginning once the reference circle was crossed. For each displacement bout, we computed the angle between the pre-movement body axis and the line to the crossed point as the movement direction (Fig. 2). Movements occurring to the left of the body axis were mirrored to the right before classification. Bouts were classified as forward (0–60° relative to the body axis), sideways (60–120°), or backward (>120°). This classification reflects three equal directional sectors in the full 360° movement space (forward/sideways/backward; 120° each), which provides a consistent reference under a null expectation of uniform movement directions. Backward movements were rare and therefore excluded from the analyses (Fig. S1). We calculated behavior using the Forward–Sideways Index (FSI): FSI = (F − S) / (F + S), where F denotes the number of forward bouts and S denotes the number of sideways bouts. FSI values range from −1 (completely sideways) to +1 (completely forward). Species with FSI < 0 were classified as sideways movers, and those with FSI > 0 as forward movers.

**Figure 2.**
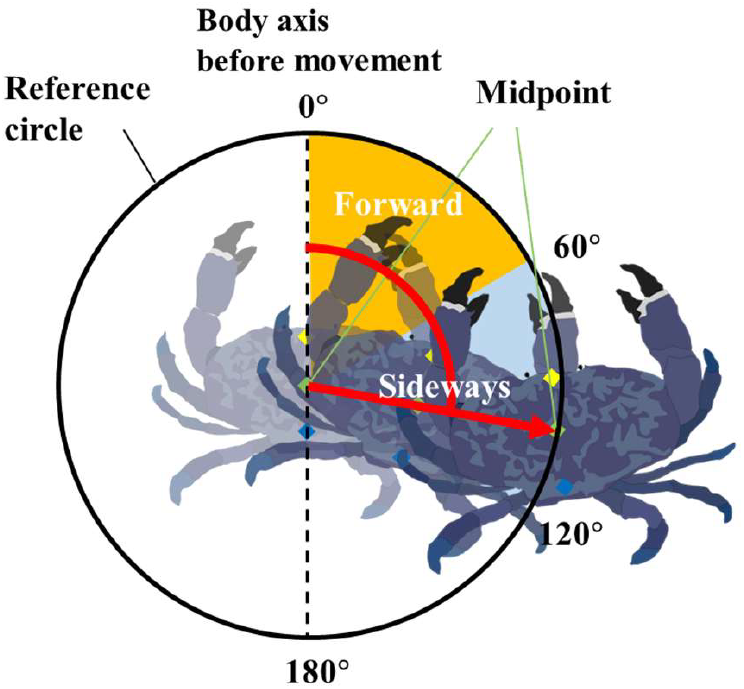
Method for determining movement directions of each crab. For each movement bout, movement direction was defined as the angle between the previous body axis (from tail to head) and the displacement vector of the body’s center (referred to as the midpoint). Displacement was measured when the midpoint reached the reference circle. These values across all movement bouts were then used to calculate the Forward–Sideways Index (FSI).

The radius of the reference circle was determined systematically for each species (Table S1). For each species, we tested circle radii from 2 mm to 200 mm in 2-mm increments and computed the FSI at each radius. When the circle is too small, digitizing noise and body wobble near the start point tend to drive FSI toward 0. When the circle is too large, animals may turn within the circle before crossing, making the estimate unstable and unreliable. To capture the biologically meaningful motions, we selected the smallest radius at which the FSI showed a clear local maximum or minimum away from zero and then remained approximately stable as the radius increased further.

### Ancestral state reconstruction

We extracted and pruned a recently published crab phylogeny (Wolfe *et al*. 2024), which was based on sequences of 10 genes (2 mitochondrial ribosomal RNA coding genes, 2 nuclear rRNA genes, and 6 nuclear protein-coding genes) and included 344 species across most major brachyuran lineages. Because our behavioral dataset did not always perfectly match the species included in (Wolfe *et al*. 2024), we reduced this tree to 44 genera, five families, and one superfamily, allowing closely related taxa to represent the observed species when the same species were unavailable. This approach enabled us to retain the terminals present in our dataset while preserving the placements of major clades relevant to this study (e.g., Eubrachyura, Raninoida).

All terminals (44 genera, five families, and one superfamily) were coded as discrete states of either forward or sideways, based on the observed FSI values (positive: forward, negative: sideways). The tree was rooted with a hermit crab terminal (Anomura), for which we assigned the observed locomotor mode of *Coenobita purpureus* from our behavioral data. The node representing the common ancestor of Brachyura and Anomura (Meiura, starting/root node) was fixed to forward as a root prior. This assumption is based on the fact that further outer groups (e.g., crayfish, Astacidea) exhibit forward locomotion (Vidal-Gadea *et al*. 2008). Ancestral states were estimated on the pruned tree using maximum likelihood under equal-rates (ER) and all-rates-different (ARD) models, with model fit compared by Akaike Information Criterion (AIC). To summarize node-wise uncertainty and transition counts, we performed stochastic character mapping (500 replicates) and reported posterior probabilities at key nodes and the posterior distributions of transitions and reversals. All analyses were conducted in R version 4.3.2 (R Core Team 2023) using the packages *ape, phytools*, and *geiger*.

### Morphological analysis

To examine the relationship between body shape and locomotor mode, we used two simple carapace shape indices: relative carapace length (CL/CW, carapace length divided by carapace width) and relative carapace depth (CD/CS, carapace depth divided by carapace size), where carapace size (CS) was defined as the geometric mean of CL, CW, and CD, where CL is carapace length, CW is carapace width, and CD is carapace depth. We used phylogenetically informed ANOVA to test whether these indices differed between forward- and sideways-moving taxa.

## Results

Of the 50 species, 35 were classified as sideways movers and 15 as forward movers based on FSI (Fig. 3; Table S1). For example, *Ranina ranina* showed an FSI of 0.89, indicating forward movement, whereas *Geothelphusa dehaani* showed an FSI of -0.70, indicating sideways movement (Fig. 4). FSI values showed a clear separation between these locomotion modes: forward movers had a median FSI of 0.82 (range: 0.24–0.94), while sideways movers had a median FSI of -0.80 (range: -1.00 to -0.39) (Fig. 3). This separation was supported by Hartigan’s dip test (*D* = 0.083, *n* = 50, *p* = 0.007), indicating significant deviation from unimodality. All species-level circular histograms are provided in Supplementary Figure S1. To further examine the underlying angle distributions, we fitted one- and two-component Gaussian mixture models to the continuous bout-angle distributions of each taxon and examined peak locations and mixture weights (Table S2). Fourteen taxa were best described by a one-component model, whereas 36 taxa were best described by a two-component model. Among the 36 taxa best described by a two-component model, 33 had a dominant component explaining at least 70% of the distribution, whereas only three taxa showed relatively balanced two-component distributions. Thus, although some taxa showed mixed directional tendencies, most taxa were dominated by a primary directional component. As an additional, data-driven check of the original FSI-based classification, we estimated a boundary from the distribution of dominant peak locations (Fig. S2; Gaussian mixture cutoff ≈ 49.4°). This peak-based classification was identical to the original FSI-based forward/sideways classification (15/35), indicating that the main classification was not dependent solely on FSI.

**Figure 3.**
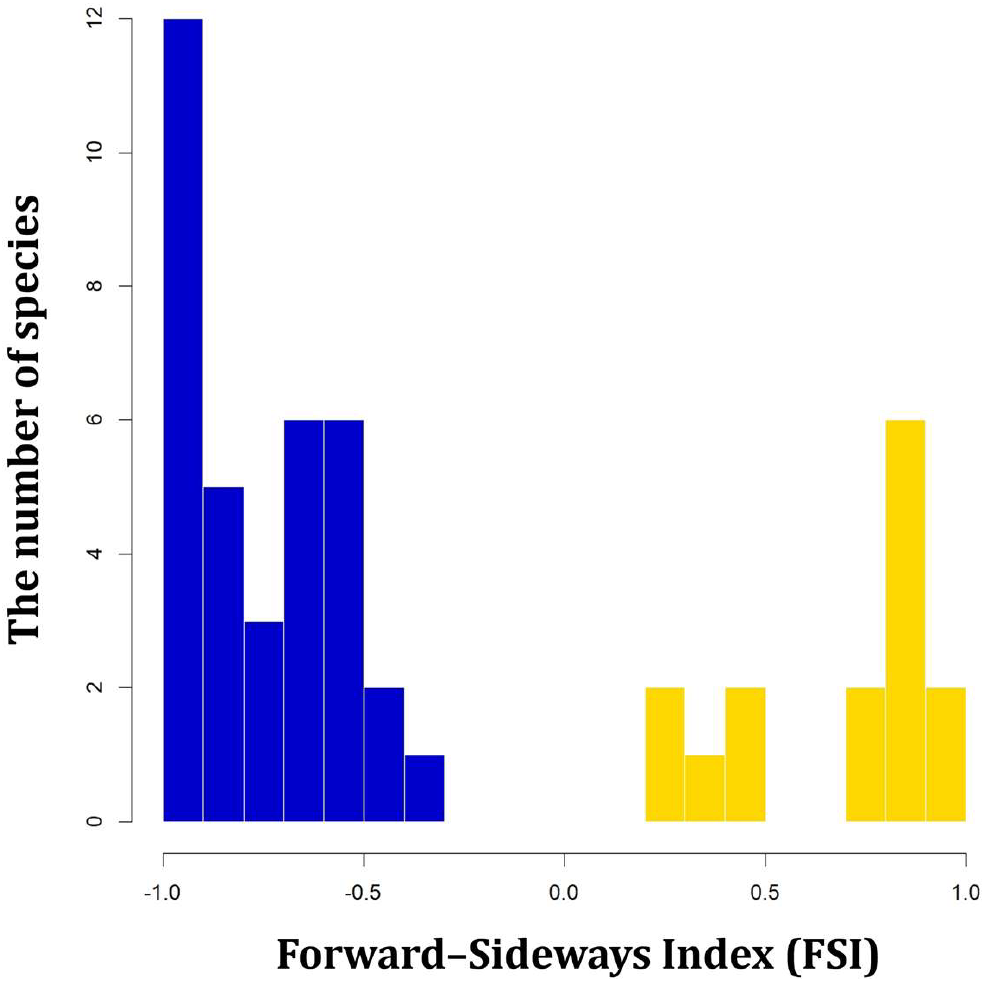
Distribution of Forward–Sideways Index (FSI) values among crab species exhibiting forward and sideways locomotion. Gold bars represent species classified as forward movers, and blue bars represent species classified as sideways movers.

**Figure 4.**
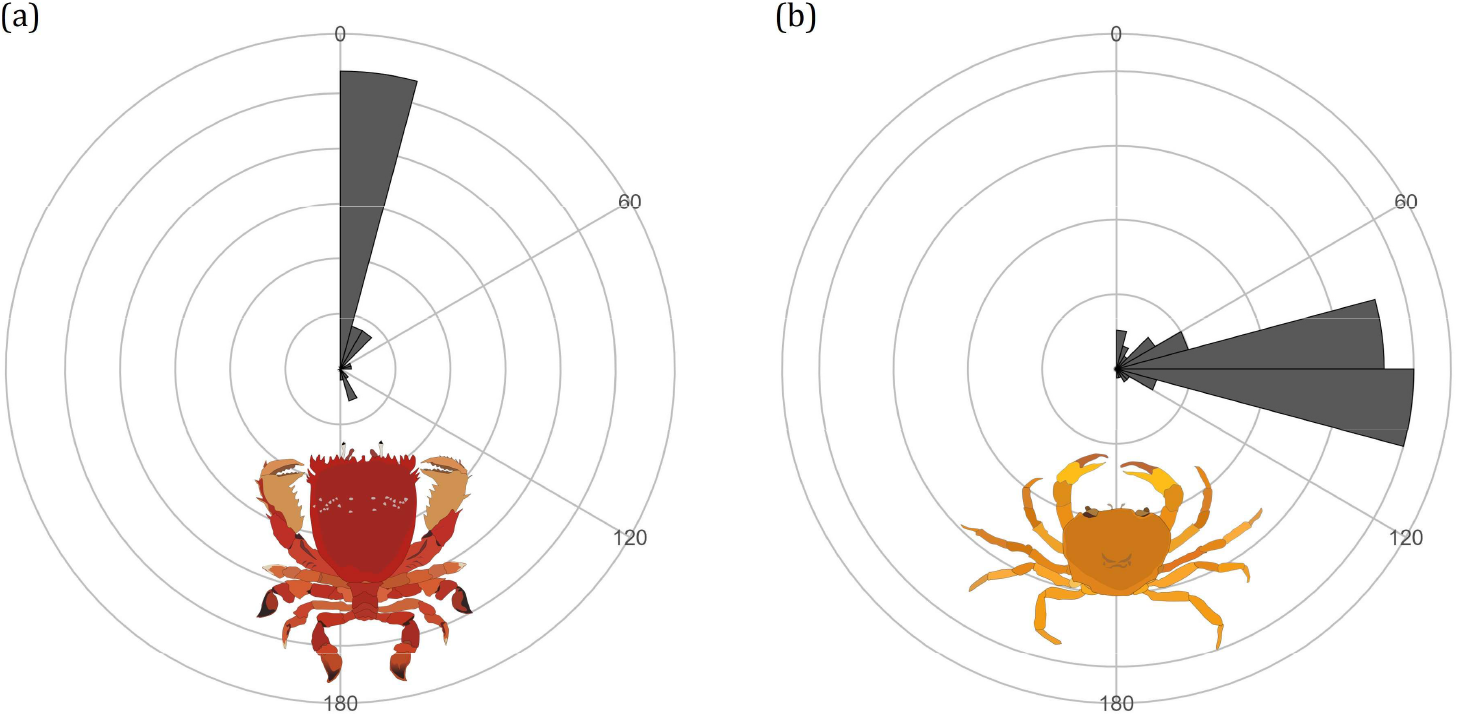
Representative circular histograms of movement directions in crabs. (a) Forward movement in *Ranina ranina* (FSI = 0.89). (b) Sideways movement in *Geothelphusa dehaani* (FSI = -0.70). The 0°-180° axis denotes the crab’s body axis before movement, with bars indicating the frequency of movement direction.

In the morphological analysis (Fig. S3), relative carapace length (CL/CW) differed significantly between forward- and sideways-moving taxa (phylogenetically informed ANOVA: *F* = 26.90, *p* < 0.001), whereas relative carapace depth (CD/CS) did not differ significantly between the two groups (*F* = 1.18, *p* = 0.403). These results suggest that locomotor mode is associated with some aspects of carapace shape, but is not explained by simple carapace flattening alone.

Model comparison favored the ARD model over the ER model (ΔAIC = 6.23; AIC weights = 0.957 vs. 0.043). Under the preferred ARD model, the transition rate from sideways to forward locomotion was estimated as 0.0029, slightly higher than the forward-to-sideways rate (0.0026). Ancestral state reconstruction placed the origin of sideways locomotion at the base of Eubrachyura (Fig. 5). Stochastic character mapping estimated the posterior probability of forward locomotion as 0.91 for the common ancestor of Brachyura, 0.79 for the common ancestor of Raninoida and Eubrachyura, and 0.24 for the eubrachyuran stem lineage. These results indicate that early-diverging lineages (i.e., Homoloida, Dromiacea, and Raninoida) retained the forward locomotion present in the common ancestor of Brachyura, and that sideways locomotion first appeared at the divergence between Raninoida and Eubrachyura. The ER model yielded a qualitatively similar placement, with only minor differences in the number of inferred transitions (Fig. S4).

**Figure 5.**
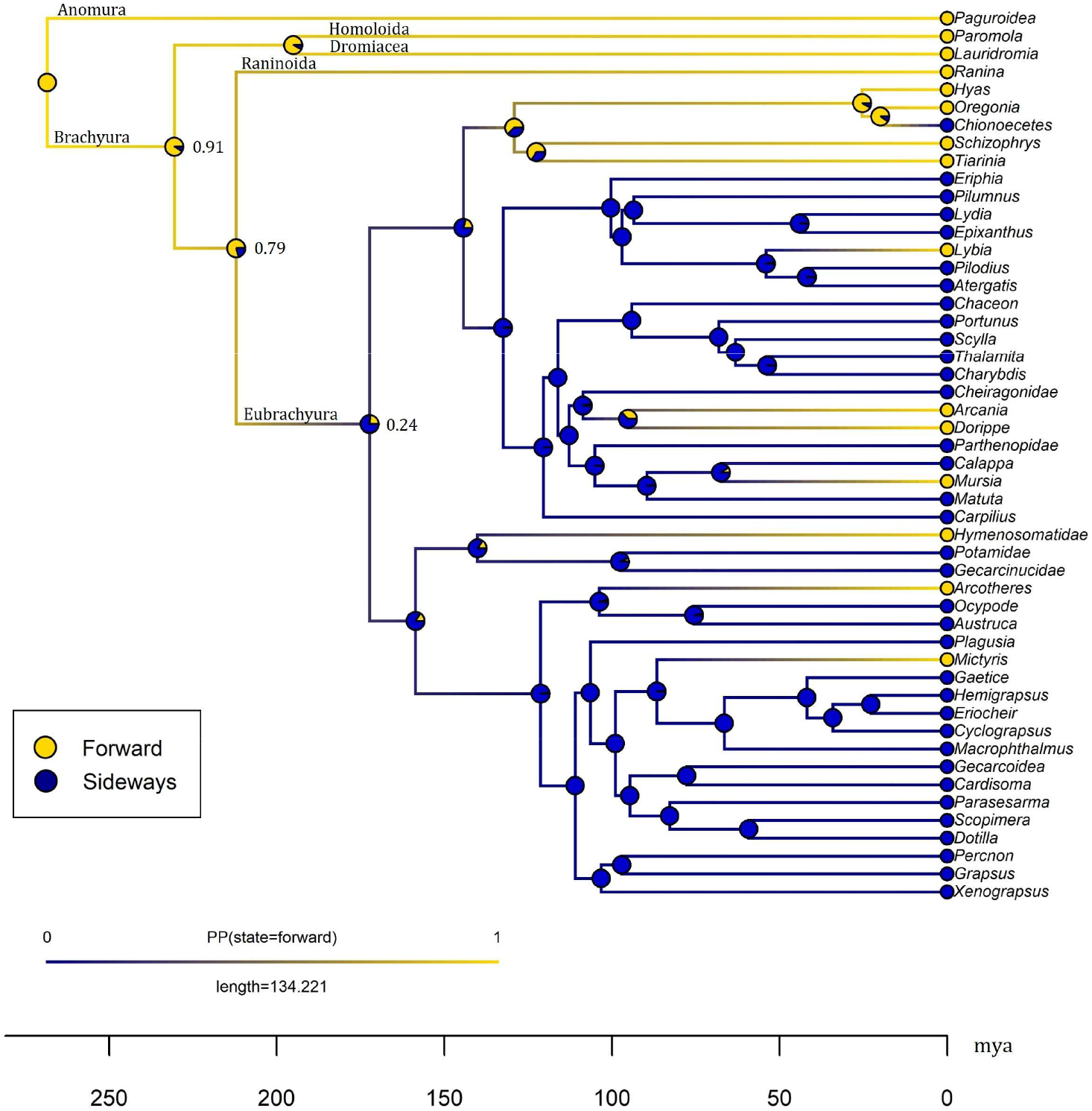
Ancestral state reconstruction of locomotion in crabs under the all-rates-different (ARD) model. Gold circles at the tips indicate forward locomotion, whereas blue circles indicate sideways locomotion. Pie charts at internal nodes and along branches represent the posterior probabilities of each locomotor state, estimated from 500 stochastic character maps. The x-axis shows geological time, scaled in millions of years before present (mya).

To place these findings in a broader phylogenetic context, we examined locomotor states alongside species richness patterns in early-diverging brachyuran lineages. Cyclodorippoida is identified as the sister group of Eubrachyura, with Raninoida forming the next outgroup (Tsang *et al*. 2011; Wolfe *et al*. 2024). Behavioral data from Raninoida indicate that these crabs move forward (Fig. 3; Table S1). In contrast, the locomotor mode of Cyclodorippidae remains unknown because these crabs inhabit deep-sea environments that preclude direct behavioral observations. Based on DecaNet (World List of Decapoda) counts of accepted extant species (DecaNet eds. 2025), Eubrachyura contains approximately 7,468 described species, whereas Cyclodorippoida includes about 110 species and Raninoida only ∼46 species. This sharp disparity highlights that the clade in which sideways locomotion is fixed—at the base of Eubrachyura or potentially in the common ancestor of Cyclodorippoida and Eubrachyura—is associated with far greater taxonomic diversity than the lineages retaining forward locomotion.

Our results also indicate that the sideways mode was largely retained across major eubrachyuran lineages. However, despite this overall stability, we inferred multiple independent reversions to forward locomotion distributed across the tree, including lineages leading to Majidae (*Hyas, Oregonia, Chionoecetes, Schizophrys, Tiarinia*), *Lybia, Arcania, Dorippe, Mursia*, Hymenosomatidae, *Arcotheres*, and *Mictyris*. Interestingly, within Majidae, *Chionoecetes* likely underwent a secondary reversion back to sideways locomotion from a forward-moving ancestor within the group. Stochastic character mapping under the ARD model estimated sideways→forward reversions at a mean of 10.2 (interquartile range, IQR = 9–12) and forward→sideways gains at a mean of 4.1 (IQR = 3–5). Under the ER model, the corresponding estimates were 10.5 (9–12) and 4.6 (3–5), respectively.

These results indicate that the evolution of sideways locomotion is rare but, once established, tends to be stably retained. Reversions to forward locomotion (and subsequent reversions back to sideways locomotion) occur under particular ecological specializations, producing a complex evolutionary pattern across true crabs.

## Discussion

Our ever-biggest behavioral dataset on crab locomotion reveals that sideways locomotion originated only once from the forward locomotion ancestor, rather than through multiple independent origins (Fig. 5). In other words, the widespread sideways locomotion across true crabs is highly conserved after being inherited from the common ancestor at the base of Eubrachyura. This suggests that modern sideways-moving crabs likely share the same anatomical, neurological, and developmental mechanisms, such as the reduction of motor neurons that control muscles of proximal legs (Vidal-Gadea & Belanger 2013). The single transition event of sideways locomotion contrasts with carcinization, which has occurred repeatedly across decapods (Tsang *et al*. 2011; Keiler *et al*. 2017; Tan *et al*. 2018; Wolfe *et al*. 2021). Carcinization has produced crab-like morphologies in several lineages of Anomura, including porcelain crabs (Porcellanidae), king crabs (Lithodidae), and the coconut crab *Birgus latro*. However, our behavioral observations indicate that porcelain crabs move predominantly backward rather than sideways, and king crabs and the coconut crab move predominantly forward (unpublished data). These examples demonstrate that even when crab-like body forms evolve, the characteristic locomotor mode (i.e., moving sideways) does not necessarily accompany them. This highlights a distinction between morphological convergence and behavioral innovation: while body forms may converge multiple times, fundamental behavioral transitions can be rare.

The species richness data show that Eubrachyura is far more diverse than its nearest outgroups, and the origin of sideways locomotion is inferred to lie around the base of the Eubrachyura. These results are consistent with the idea that this unique locomotor mode acted as a key innovation that contributed to the evolutionary diversification of Eubrachyura. Note, however, that a key innovation is not the only process driving adaptive radiation, and the innovation may not always result in radiation (Fürsich & Jablonski 1984; Miller *et al*. 2023). External factors, such as ecological opportunity provided by mass extinction, are also critical for evolutionary diversification (Stroud & Losos 2016). Based on the divergence times reported by (Wolfe *et al*. 2024), the origin of sideways locomotion falls around ∼200 Mya (earliest Jurassic, immediately post–Triassic–Jurassic extinction), a recovery interval marked by Pangaean rifting, expansion of shallow-marine habitats, and the early Mesozoic Marine Revolution—conditions that typically increase ecological opportunity (Buatois *et al*. 2016; Schoepfer *et al*. 2022). Disentangling the relative roles of intrinsic innovation and extrinsic environmental change will require trait-dependent diversification analyses (Maddison *et al*. 2007), fossil-informed timelines, and performance tests that link sideways movement to adaptive advantages.

Sideways locomotion may confer several adaptive advantages. One likely advantage is the ability to move rapidly at similar speeds in both lateral directions (Vidal-Gadea *et al*. 2008; Wolfe *et al*. 2021), which is also supported by an experiment using crab-like robots (Chen *et al*. 2022). Having multiple locomotor directions is highly advantageous for escaping from predators, not only by making the escape direction unpredictable but also by providing multiple optimal escape routes (Domenici *et al*. 2011; Kawabata *et al*. 2023). Other possible advantages may include movement through confined spaces and visual-field sampling during locomotion, although these possibilities remain untested and will require future biomechanical and sensory studies. An alternative explanation is that sideways locomotion is a secondary phenomenon resulting from a flattened body structure that limits the distance between the legs and thereby makes forward movement difficult. However, our additional morphological analysis showed that relative carapace depth (CD/CS) did not differ significantly between forward- and sideways-moving taxa, whereas relative carapace length (CL/CW) differed significantly. Therefore, sideways locomotion cannot be explained solely by carapace flattening, and the morphological correlates of sideways locomotion are more complex.

Despite these potential adaptive advantages, sideways locomotion appears to have evolved rarely across the animal kingdom; it has occurred only in true crabs (Fig. 5), and potentially also in crab spiders (Wilcox 2017) and leafhopper nymphs (Chasen *et al*. 2014). One possible reason is that this locomotor mode fundamentally changes the behavioral axis, potentially affecting a wide range of behaviors such as predator avoidance, shelter use, burrowing, mating, and foraging (Atkinson & Eastman 2015; Crane 2015; Asakura 2016; Takeshita & Nishiumi 2022). In this sense, the origin of sideways locomotion may represent a rare and major behavioral transition rather than a simple change in movement direction alone.

Nevertheless, sideways locomotion does not appear to be universally favored across all ecological contexts. After the transition to sideways locomotion, crabs have experienced at least six independent reversions to forward locomotion (Fig. 5). These reversions are particularly associated with major changes in life history traits. For example, soldier crabs (*Mictyris*) predominantly use forward walking (Fig. S1; Table S1) in a way biomechanically similar to other forward-walking animals rather than sideways-walking crabs (Sleinis & Silvey 1980). Soldier crabs are unique for their gregarious nature and coordinated collective movements (Murakami *et al*. 2014), which may have brought them back to forward locomotion. Similarly, majoid crabs (e.g., *Oregonia, Tirarinia*) camouflage themselves with seaweed (Sato & Wada 2000; Hultgren & Stachowicz 2009), and pea crabs (e.g., *Arcotheres*) live hidden inside bivalves and other invertebrates, relying on their hosts for protection (de Gier & Becker 2020). Given that the major benefit of sideways locomotion is rapid escape from predators (Vidal-Gadea *et al*. 2008; Wolfe *et al*. 2021), these examples with alternative strategies of predator avoidance may no longer need sideways locomotion, resulting in secondary losses. These exceptional forward-moving species imply that it is costly to maintain sideways locomotion in crabs, and that they retain evolutionary flexibility to lose this locomotor mode under certain ecological pressures.

Modes of locomotion—such as walking, swimming, and flying—fundamentally shape how animals interact with the environment, affecting behaviors related to foraging, predator avoidance, and reproduction (Alexander 2003; Domenici 2010). Sideways locomotion in true crabs is also a critical change in the mode of locomotion, whose evolutionary history is characterized by rarity, stability, and occasional reversions. Our case study illustrates how major innovations can open new adaptive opportunities, yet remain constrained by both phylogenetic history and ecological context. By integrating direct behavioral observations with a robust phylogenetic framework, this study expands our understanding of how animal locomotor modes diversify and persist through evolutionary time.

## Acknowledgements

We sincerely thank the staff and the students of the Kawabata Laboratory, Nagasaki University, and the Huang Laboratory, National Kaohsiung University of Science and Technology, for their assistance with crab sampling, experiments, and video analysis. We are deeply grateful to the staff of the Susami Crustacean Aquarium and the Wakayama Prefectural Museum of Natural History for generously providing experimental animals, assisting with field sampling, and allowing us to use their facilities for behavioral observations. We sincerely thank Takehiro Sato and the staff of the Kanagawa Prefectural Museum of Natural History for their generous assistance in measuring crab morphology. We are also grateful to Akinori Yamada, Nagasaki University, for his assistance with the preliminary analysis of crab phylogeny construction using genomic data.

## References

Alexander, R.M. (2003). Principles of animal locomotion. Princeton University Press, New Jersey, USA.

Asakura, A. (2016). The evolution of mating systems in decapod crustaceans. In: Decapod crustacean phylogenetics. CRC Press, pp. 133–194.

Astudillo-Clavijo, V., Arbour, J.H. & López-Fernández, H. (2015). Selection towards different adaptive optima drove the early diversification of locomotor phenotypes in the radiation of Neotropical geophagine cichlids. BMC Evol. Biol., 15, 77.

Atkinson, R.J.A. & Eastman, L.B. (2015). Burrow dwelling in Crustacea. The natural history of the Crustacea, 2, 78–117.

Buatois, L.A., Carmona, N.B., Curran, H.A., Netto, R.G., Mángano, M.G. & Wetzel, A. (2016). The Mesozoic Marine Revolution. In: The Trace-Fossil Record of Major Evolutionary Events: Volume 2: Mesozoic and Cenozoic (eds. Mángano, MG & Buatois, LA). Springer Netherlands Dordrecht, pp. 19–134.

Burress, E.D. & Wainwright, P.C. (2019). Adaptive radiation in labrid fishes: A central role for functional innovations during 65 My of relentless diversification. Evolution, 73, 346–359.

Chasen, E.M., Dietrich, C., Backus, E.A. & Cullen, E.M. (2014). Potato leafhopper (Hemiptera: Cicadellidae) ecology and integrated pest management focused on Alfalfa. Journal of Integrated Pest Management, 5, A1–A8.

Chen, Y., Grezmak, J.E., Graf, N.M. & Daltorio, K.A. (2022). Sideways crab-walking is faster and more efficient than forward walking for a hexapod robot. Bioinspir Biomim, 17.

Crane, J. (2015). Fiddler crabs of the world: Ocypodidae: genus Uca. Princeton University Press.

de Gier, W. & Becker, C. (2020). A review of the ecomorphology of pinnotherine pea crabs (Brachyura: Pinnotheridae), with an updated list of symbiont-host associations. Diversity, 12, 431.

DecaNet eds. (2025). DecaNet. Accessed at https://www.decanet.info on 2025-09-16.

Domenici, P. (2010). Fish locomotion: an eco-ethological perspective. CRC Press.

Domenici, P., Blagburn, J.M. & Bacon, J.P. (2011). Animal escapology I: Theoretical issues and emerging trends in escape trajectories. J. Exp. Biol., 214, 2463–2473.

Fürsich, F.T. & Jablonski, D. (1984). Late triassic naticid drillholes: Carnivorous gastropods gain a major adaptation but fail to radiate. Science, 224, 78–80.

Faulkes, Z. (2006). The locomotor toolbox of the spanner crab, Ranina ranina (Brachyura, Raninidae). Crustaceana, 143–155.

Feiner, N., Jackson, I.S.C., Stanley, E.L. & Uller, T. (2021). Evolution of the locomotor skeleton in Anolis lizards reflects the interplay between ecological opportunity and phylogenetic inertia. Nature Communications, 12, 1525.

Grimaldi, D. & Engel, M.S. (2005). Evolution of the Insects. Cambridge University Press.

Hedrick, B.P., Dickson, B.V., Dumont, E.R. & Pierce, S.E. (2020). The evolutionary diversity of locomotor innovation in rodents is not linked to proximal limb morphology. Sci. Rep., 10, 717.

Higham, T.E., Birn-Jeffery, A.V., Collins, C.E., Hulsey, C.D. & Russell, A.P. (2015). Adaptive simplification and the evolution of gecko locomotion: Morphological and biomechanical consequences of losing adhesion. Proceedings of the National Academy of Sciences, 112, 809–814.

Hultgren, K. & Stachowicz, J. (2009). Evolution of decoration in majoid crabs: a comparative phylogenetic analysis of the role of body size and alternative defensive strategies. The American Naturalist, 173, 566–578.

Kawabata, Y., Akada, H., Shimatani, K.I., Nishihara, G.N., Kimura, H., Nishiumi, N. et al. (2023). Multiple preferred escape trajectories are explained by a geometric model incorporating prey’s turn and predator attack endpoint. eLife, 12, e77699.

Keiler, J., Wirkner, C.S. & Richter, S. (2017). One hundred years of carcinization – the evolution of the crab-like habitus in Anomura (Arthropoda: Crustacea). Biol. J. Linn. Soc., 121, 200–222.

Maddison, W.P., Midford, P.E. & Otto, S.P. (2007). Estimating a binary character’s effect on speciation and extinction. Syst. Biol., 56, 701–710.

Miller, A.H. (1949). Some ecologic and morphologic considerations in the evolution of higher taxonomic categories. In: Ornithologie als biologische Wissenschaft, p. 84.

Miller, A.H., Stroud, J.T. & Losos, J.B. (2023). The ecology and evolution of key innovations. Trends Ecol. Evol., 38, 122–131.

Morrison, C.L., Harvey, A.W., Lavery, S., Tieu, K., Huang, Y. & Cunningham, C.W. (2002).Mitochondrial gene rearrangements confirm the parallel evolution of the crab-like form. Proc. R. Soc. Lond. B, 269, 345–350.

Murakami, H., Tomaru, T., Nishiyama, Y., Moriyama, T., Niizato, T. & Gunji, Y.-P. (2014). Emergent runaway into an avoidance area in a swarm of soldier crabs. PloS one, 9, e97870.

R Core Team (2023). R: The R project for statistical computing.

Sato, M. & Wada, K. (2000). Resource utilization for decorating in three intertidal majid crabs (Brachyura: Majidae). Mar. Biol., 137, 705–714.

Schoepfer, S.D., Algeo, T.J., van de Schootbrugge, B. & Whiteside, J.H. (2022). The Triassic– Jurassic transition – A review of environmental change at the dawn of modern life. Earth-Sci. Rev., 232, 104099.

Sleinis, S. & Silvey, G.E. (1980). Locomotion in a forward walking crab. Journal of comparative physiology, 136, 301–312.

Stroud, J.T. & Losos, J.B. (2016). Ecological opportunity and adaptive radiation. Annual Review of Ecology, Evolution, and Systematics, 47, 507–532.

Takeshita, F. & Nishiumi, N. (2022). Social behaviors elevate predation risk in fiddler crabs: quantitative evidence from field observations. Behav. Ecol. Sociobiol., 76, 162.

Tan, M.H., Gan, H.M., Lee, Y.P., Linton, S., Grandjean, F., Bartholomei-Santos, M.L. et al. (2018). ORDER within the chaos: Insights into phylogenetic relationships within the Anomura (Crustacea: Decapoda) from mitochondrial sequences and gene order rearrangements. Mol. Phylogenet. Evol., 127, 320–331.

Tsang, L.M., Chan, T.Y., Ahyong, S.T. & Chu, K.H. (2011). Hermit to king, or hermit to all: multiple transitions to crab-like forms from hermit crab ancestors. Syst. Biol., 60, 616–629.

Vidal-Gadea, A.G. & Belanger, J.H. (2013). The evolutionary transition to sideways-walking gaits in brachyurans was accompanied by a reduction in the number of motor neurons innervating proximal leg musculature. Arthropod Struct. Dev., 42, 443–454.

Vidal-Gadea, A.G., Rinehart, M.D. & Belanger, J.H. (2008). Skeletal adaptations for forwards and sideways walking in three species of decapod crustaceans. Arthropod Struct. Dev., 37, 95–108.

Wilcox, J.T. (2017). Crab spider: grasslands predator hiding in plain sight. California Native Grasslands Association, pp. 3–4.

Wolfe, J.M., Ballou, L., Luque, J., Watson-Zink, V.M., Ahyong, S.T., Barido-Sottani, J. et al. (2024). Convergent adaptation of true crabs (Decapoda: Brachyura) to a gradient of terrestrial environments. Syst. Biol., 73, 247–262.

Wolfe, J.M., Luque, J. & Bracken-Grissom, H.D. (2021). How to become a crab: Phenotypic constraints on a recurring body plan. BioEssays, e2100020.

