## Supplementary Figures for "Evolution of sideways locomotion in crabs"

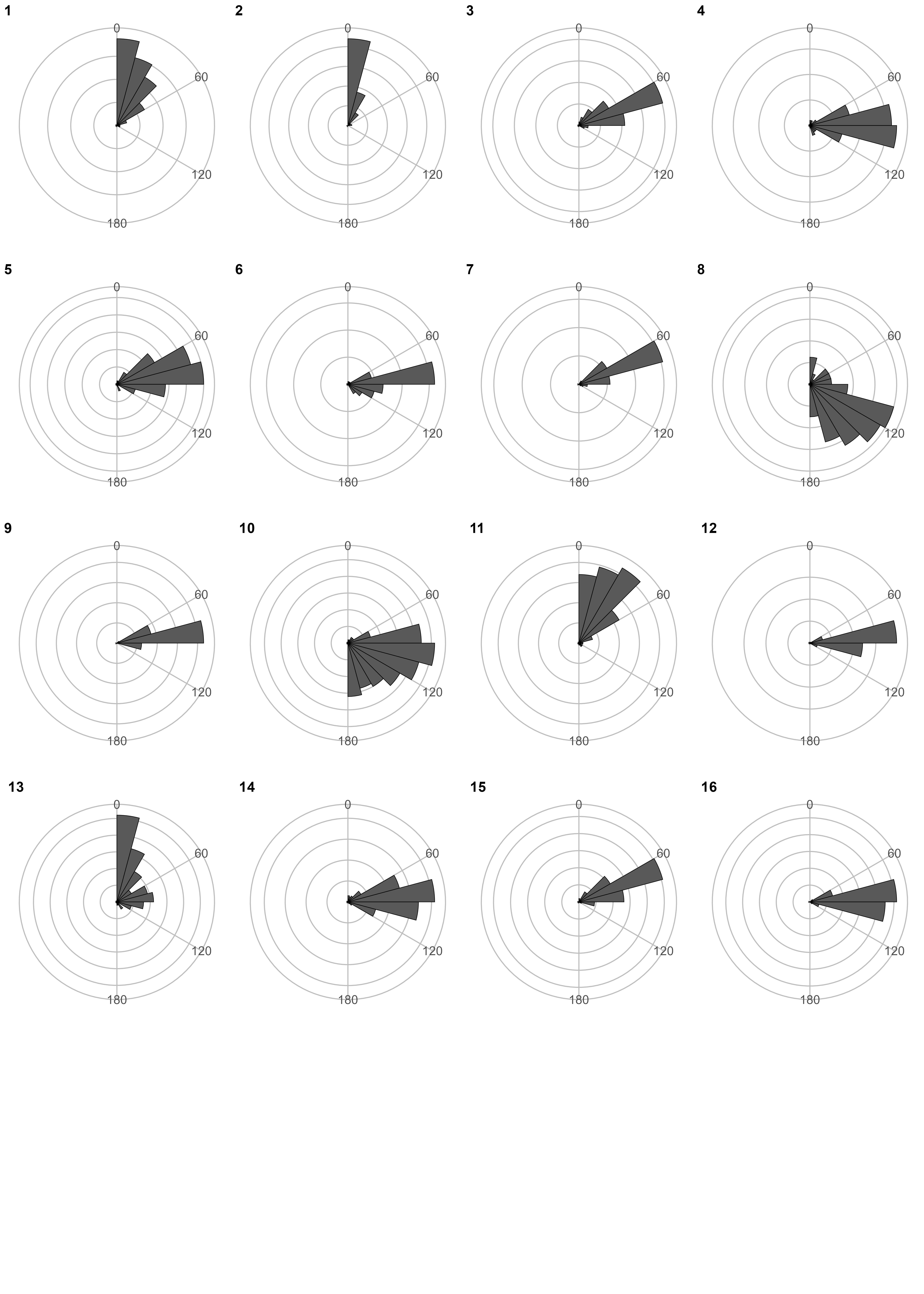

**Figure S1.** Circular histograms showing movement direction for every crab on our dataset. 1. *Arcania heptacantha* 2. *Arcotheres sinensis* 3. *Atergatis floridus* 4. *Austruca lactea* 5. *Calappa philargius* 6. *Cardisoma carnifex* 7. *Carpilius convexus* 8. *Chaceon granulatus* 9. *Charybdis (Charybdis) japonica* 10. *Chionoecetes opilio* 11. *Coenobita purpureus* 12. *Cyclograpsus intermedius* 13. *Dorippe sinica* 14. *Dotilla wichmanni* 15. *Enoplolambrus validus* 16. *Epixanthus frontalis*

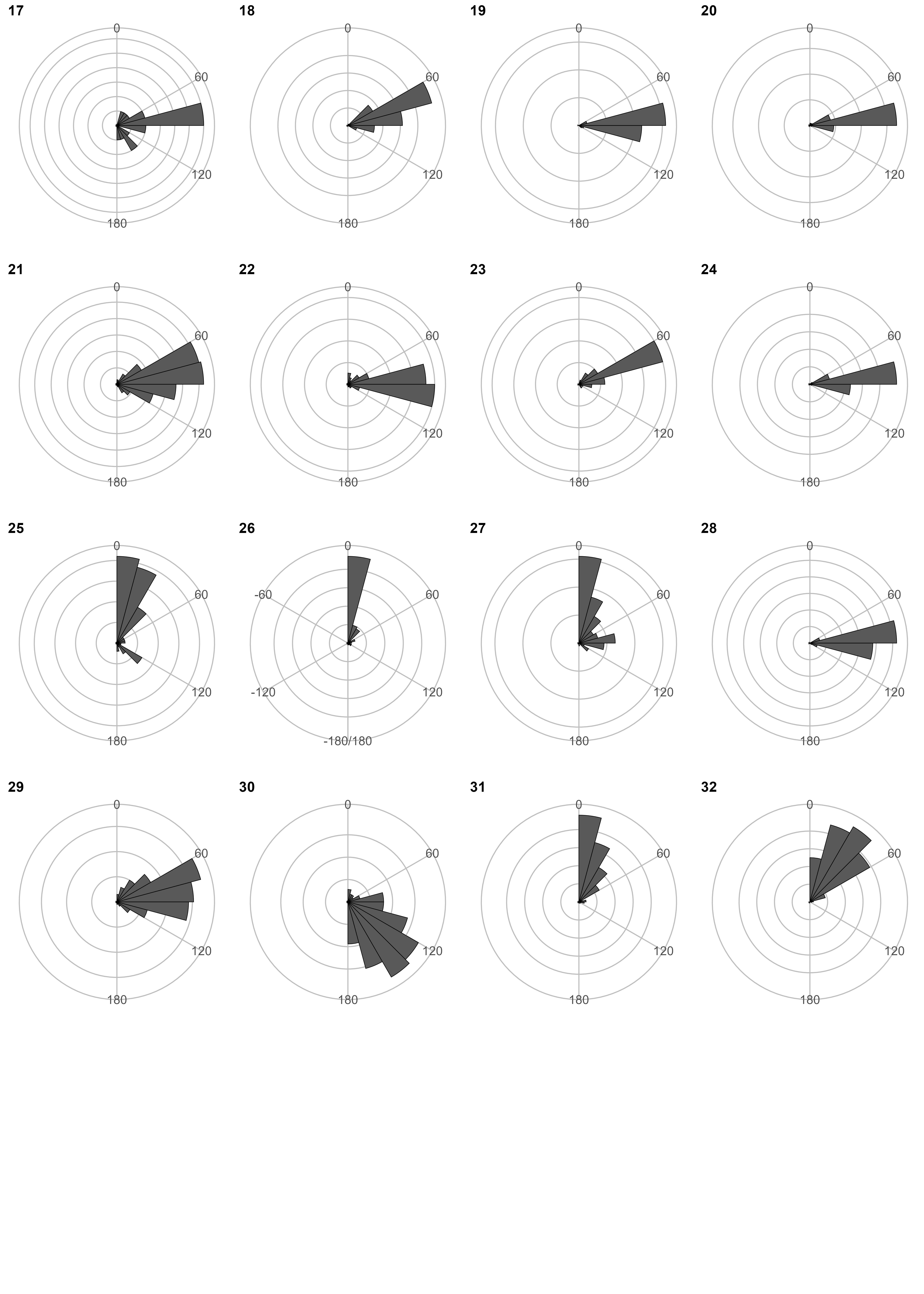

**Figure S1.** (Continued) 17. *Erimacrus isenbeckii* 18. *Eriocheir japonica* 19. *Eriphia ferox* 20. *Gaetice depressus* 21. *Gecarcoidea lalandii* 22. *Geothelphusa dehaani* 23. *Grapsus albolineatus* 24. *Hemigrapsus sanguineus* 25. *Hyas alutaceus* 26. *Lauridromia dehaani* 27. *Lybia tessellate* 28. *Lydia annulipes* 29. *Macrophthalmus (Mareotis) japonicus* 30. *Matuta victor* 31. *Mictyris brevidactylus* 32. *Mursia armata*

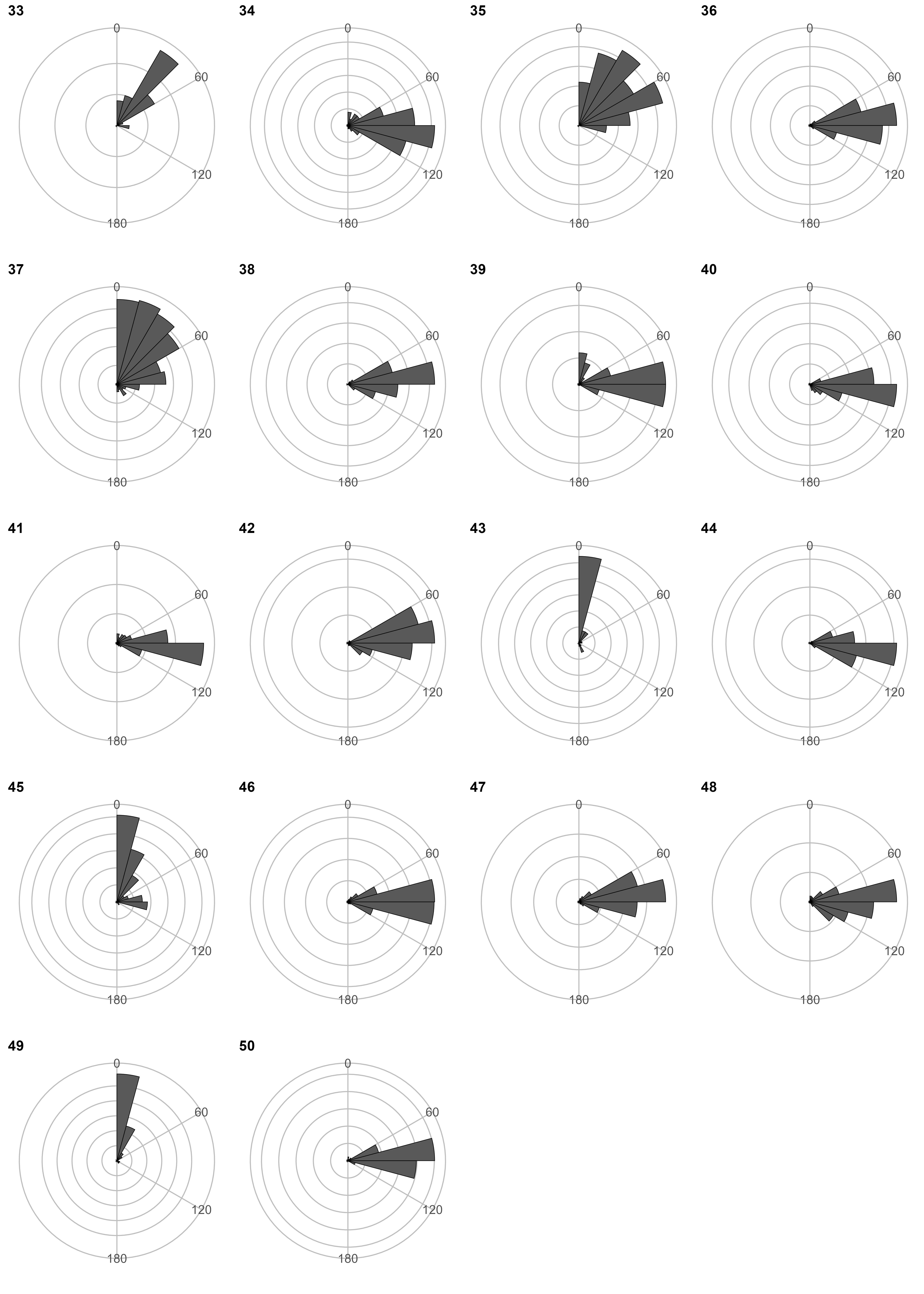

**Figure S1.** (Continued) 33. *Neohymenicus orientalis* 34. *Ocypode stimpsoni* 35. *Oregonia gracilis* 36. *Parasesarma pictum* 37. *Paromola japonica* 38. *Percnon planissimum* 39. *Pilodius areolatus* 40. *Pilumnus vespertilio* 41. *Plagusia squamosa* 42. *Portunus pelagicus* 43. *Ranina ranina* 44. *Sayamia germaini* 45. *Schizophrys aspera* 46. *Scopimera globosa* 47. *Scylla serrata* 48. *Thalamita sima* 49. *Tiarinia cornigera* 50. *Xenograpsus testudinatus*

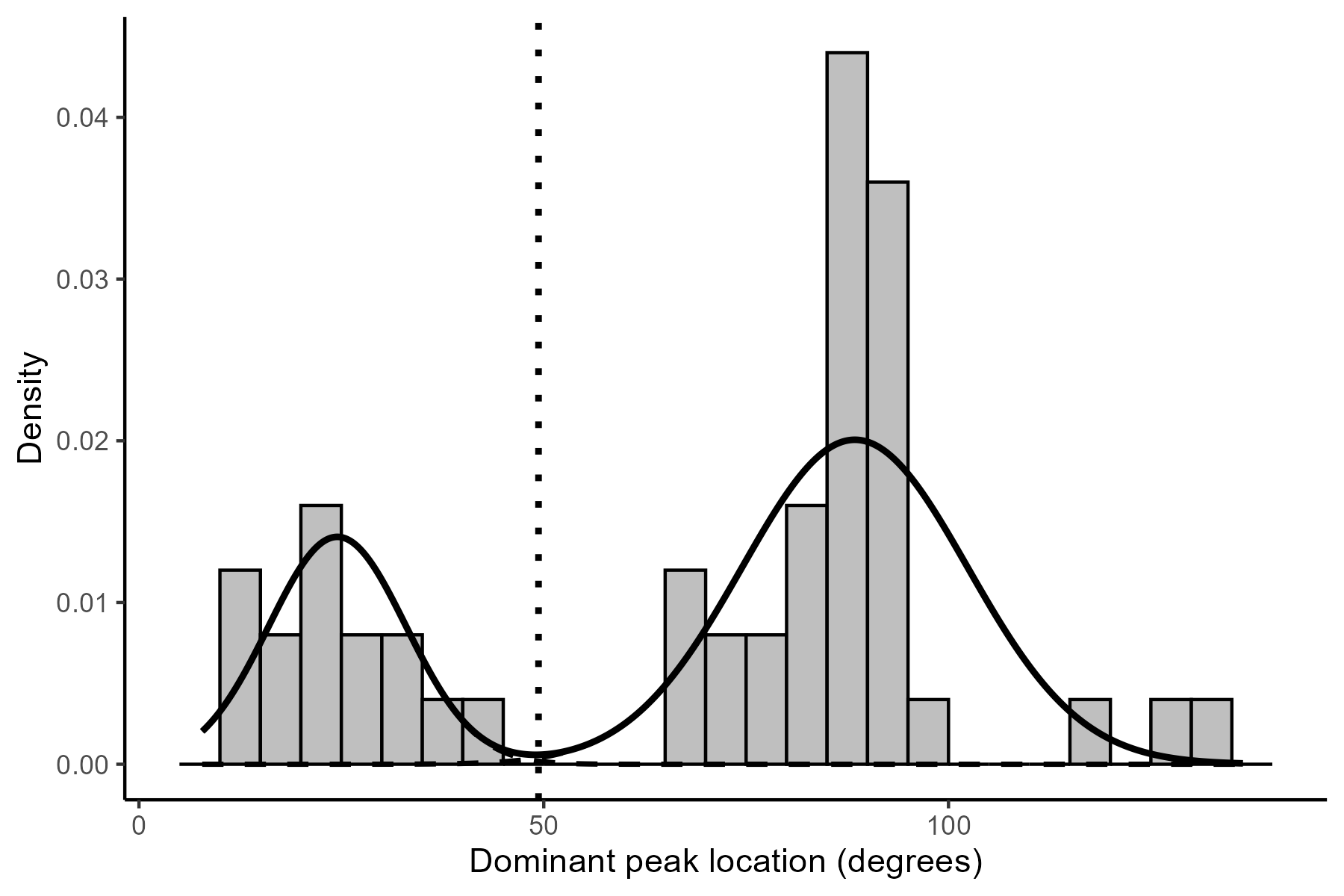

**Figure S2.** Gaussian mixture fit to dominant peak locations. Dominant peak locations were extracted from mixture models fitted to each taxon’s continuous angle distribution (Table S2). A two-component Gaussian mixture model fitted to these locations yielded an estimated cutoff at 49.4° (dotted line). Classifying taxa using this data-informed cutoff produced an identical forward/sideways classification to our original approach using Forward-Sideways index (15 forward, 35 sideways).

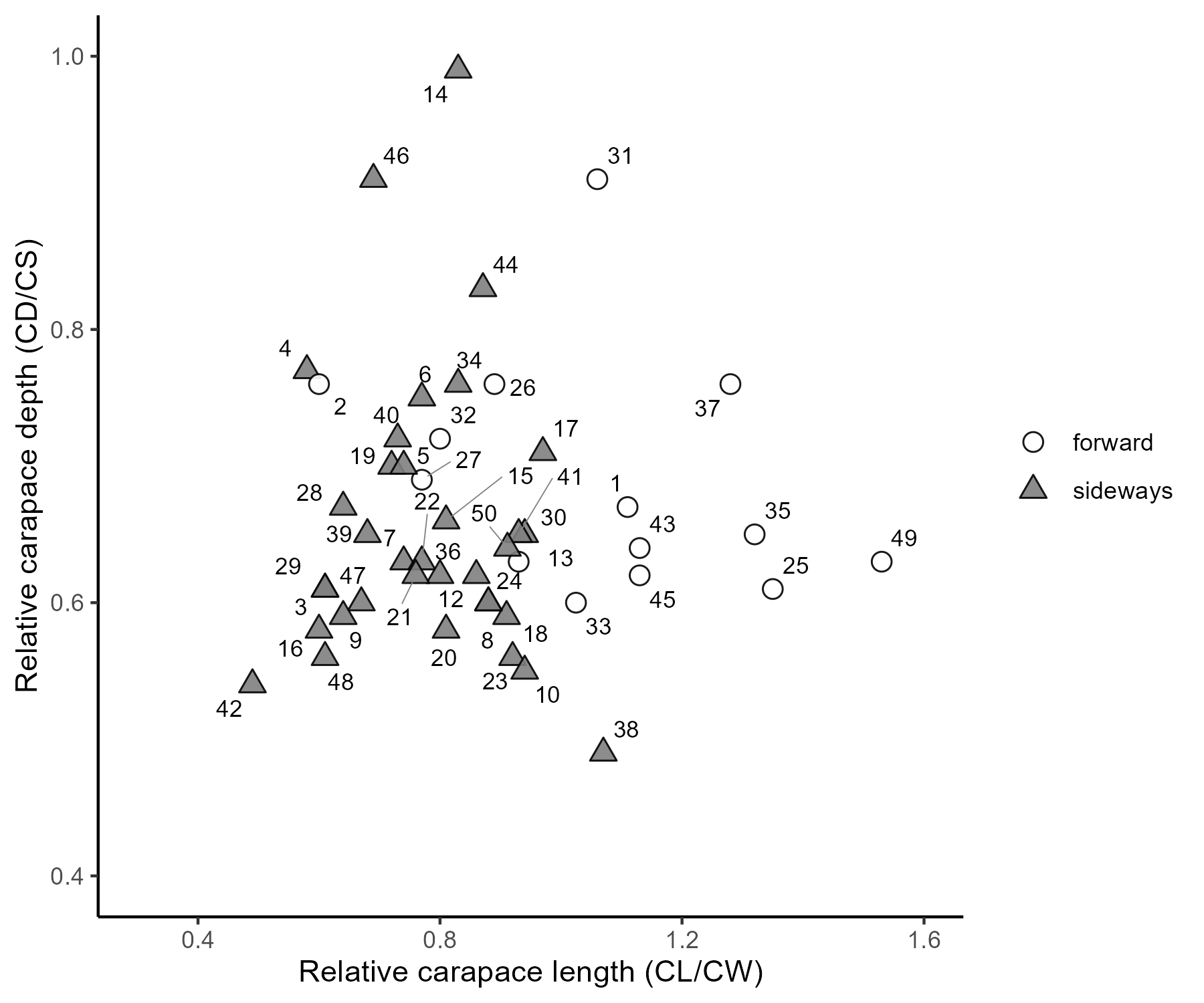

**Figure S3.** Morphospace of crab locomotor modes based on relative carapace length (CL/CW) and relative carapace depth (CD/CS). Forward- and sideways-moving taxa overlapped broadly in morphospace, although phylogenetically informed ANOVA detected a significant difference in CL/CW but not in CD/CS. Numbers in the plot correspond to the species numbers shown in Figure S1.

CL, carapace length; CW, carapace width; CD, carapace depth; CS, carapace size, defined as the geometric mean of CL, CW, and CD.

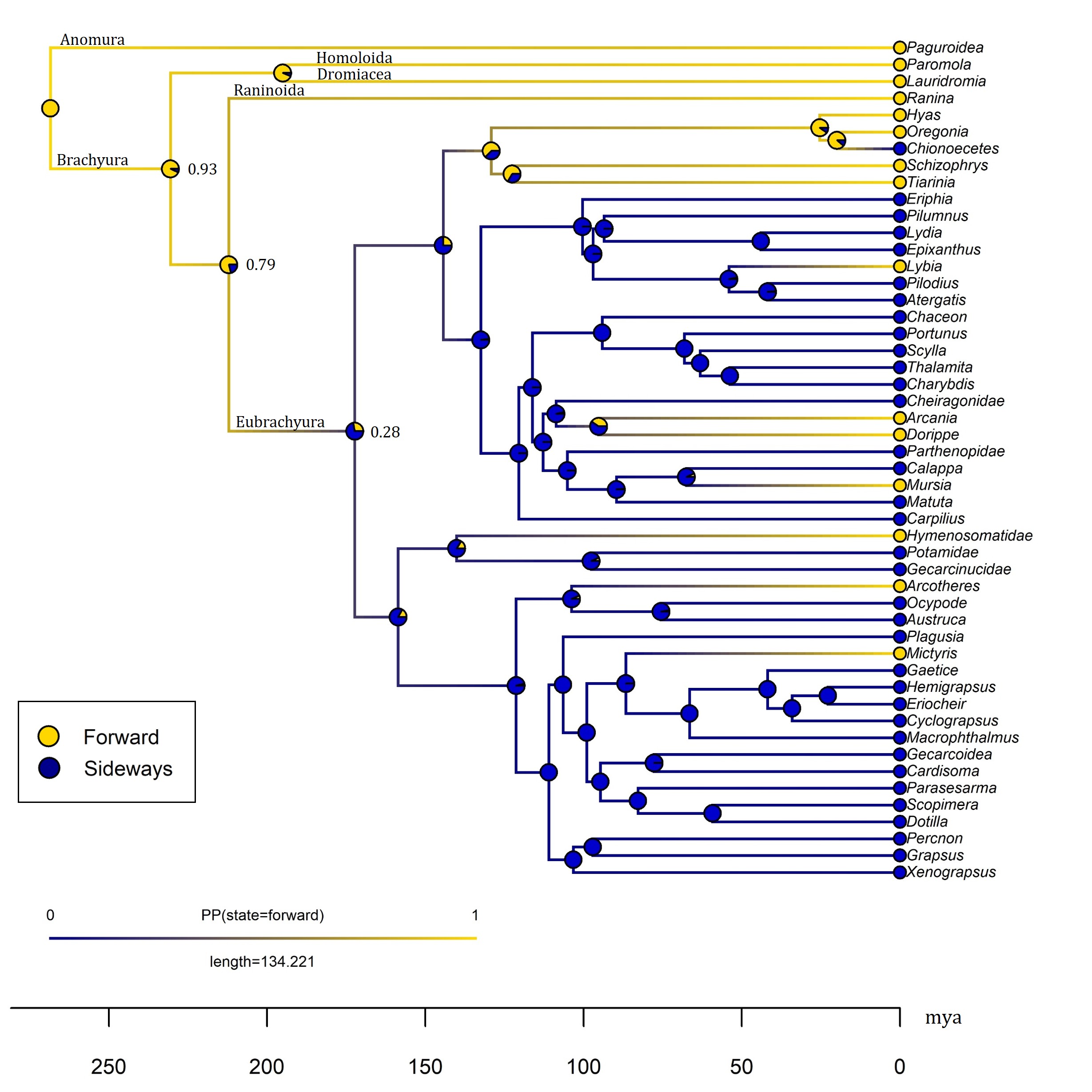

**Figure S4.** Ancestral state reconstruction of locomotion in crabs under the equal-rates (ER) model. Gold circles at the tips indicate forward locomotion, whereas blue circles indicate sideways locomotion. Pie charts at internal nodes and along branches represent the posterior probabilities of each locomotor state, estimated from 500 stochastic character maps. The x-axis shows geological time, scaled in millions of years before present (mya).
